# Dietary sulfate-driven and gut dysbiosis-triggered breast cancer-related gene upregulation

**DOI:** 10.1101/440578

**Authors:** CHEN YanPingf, LIAO Tao, TAN LiLi, CHEN DongMei, XU Qin, Song JianPing, ZHEN QingPing

## Abstract

By gut microbiota metagenomic analysis, we found that the abundance of sulfatase-secreting bacteria (SSB) in the gut of mice fed chondroitin sulfate (CS) increases with significant individual difference. The fluctuation of lipopolysaccharide (LPS) and pro-inflammatory indicators with significant individual and tissue variations was also observed. After mice were fed mixed with CS or injected separately with LPS, the breast cancer-related transcriptional factor genes, *BCL11A* and *RUNX1*, were upregulated, whereas the tumor suppressor gene, *TP53BP1*, were downregulated. Further, the mammary myopithelium marker CK5/6, the mammary hyperplasia marker Ki-67, and other tumor markers were also upregulated. While the exogenous estradiol does not induce the expression of *BCL11A*, *RUNX1*, and *TP53BP1*, the estrogen receptor (ER) agonist Fulvestrant that mimics estradiol action not only elevates estradiol concentrations, but also upregulates tumor marker expression levels, revealing that ER inflammatory inactivation and hyperestrogenemia induction might be the etiological cues of breast cancer origin. This study has preliminarily established a possible correlation of gut microbiota dysbiosis and chronic low-grade inflammation with the early-phase onset of breast cancer in mice. The statistical insignificance of test data was attributed to the individual difference of gut microbiota compositions, which determining the individual and tissue variations of systemic inflammation.

---

The statistical data in 2015 have indicated that the global incidence rate of breast cancer was up to 12% among women^[1]^. In all patients, only 2-3% cases are belonging to familiar early-onset breast cancer, which are resulted from *BRCA1* or *BRCA2* gene mutation, and others are non-genetically late-onset breast cancer^[2]^, whose etiological reasons remain unknown. Nevertheless, accumulating evidence shows that hyperestrogenemia has been considered as the critical inducer of breast cancer^[3]^, in a possible mechanism underlying that estrogens stimulate estrogen receptors (ERs) and further activate the downstream cell division-promoting transcription^[4]^. For the source of high-level estrogens, the current explanation is that colon cell-secreted β-glucuronidase converts estrogens from the bound-types to the free types, enabling re-adsorption via the enterohepatic circulation^[5]^.

On the other hand, some evidence indicates that the unhealthy lifestyles, including a high-fat diet^[6]^, smoking^[7]^, drunk^[8]^, are significantly associated with breast cancer incidence, and a possible link of gut microbiota disorder to breast cancer incidence has been also concerned^[9]^. According to these cues, we suggest here a hypothesis of the “gut origin of breast cancer”, which addresses that gut dysbiosis from opportunistic infection leads to colon mucosal damaged, bacterial endotoxin leaked into blood circulation, and systemic inflammation induced^[10]^. Along with the inflammatory lesion and functional inactivation of ERs, estrogen levels should be complementarily elevated, and oncogenes accordingly upregulated or tumor suppressors downregulated, eventually inducing mammary gland hyperplasia (benign tumor) or transformation (malignant cancer).

Because animal-derived chondroitin sulfate (CS) was proven to trigger opportunistic infection from the overgrowth of sufatase-secreting bacteria (SSB) and sulfate-reducing bacteria^[11]^, we fed CS and SSB to female mice to mimic gut dysbiosis and induce chronic low-grade inflammation^[12]^. A possibility of the interaction of CS-rich meat diets with gut microbiota communities raise a breast cancer risk was evaluated by determining the expression levels of mammary tumor-related transcription factors and tumor markers. A cause of hyperestrogenemia during breast cancer incidence was proposed by comparing the expression levels of mammary tumor-related transcriptional factors upon injection of the exogenous estrogen, estradiol (ED)^[13]^, or the ERs agonist, Fulvestrant (FS)^[14]^. An association of the individual gene expression difference with the inconsistent experimental repeatability was revealed by the metagenomic gut microbiome analysis^[15]^.

The present study preliminarily dissected the putative mechanism underlying that gut dysbiosis induces mammary tumor by eliciting systemic inflammation. On this basis, it should be realistic to improve gut ecosystems, eradicate tumorigenic cues, and decline carcinogenic risks by artificial attempts, which would create an innovative era for the early-phase intervention of gut dysbiosis and effective prevention of breast cancer.

## 1 Materials and Methods

### 1.1 Animal group and reagent treatment

The specific pathogen free BALB/c mice (female, 18-22 g) and Kunming mice (female, 30±2 g) were provided by the experimental animal center in Guangzhou University of Chinese Medicine. The animals were allowed feeding routinely and drinking freely, and divided randomly into 6 groups with 2-4 mice in each group. ➀ Group *ad libitum* (AL); ➁ Group CS (FocusChem, Shandong, China); ➂Group CS+*Bacillus cereus* (BC, Huankai Microb, Guangdong, China); ➃Group CS+BC+FS(MCE); ➄Group LPS (Sigma); ➅Group low-dose ED (Alfa Aesar); ➆ Group high-dose ED.

0.25 ml CS (1.0 g/kg) was used to intragastrical feed group ➁ mice, and repeat after 2w; A solution containing 10^4^ BC was used to intragastrical feed group ➂ mice. On the next day, they were intragastrically fed by 0.25 ml CS (1.0 g/kg), and repeat after 2w; FS (500 μg/kg) was used to peritoneal inject group ➃ mice, 1 time/d for 60 d; LPS (0.25 mg/kg) was used to peritoneal inject group ➄ mice, 1 time every other day for 60 d; 0.5 mg/kg and 4.0 mg/kg ED were used to inject group ➅ and ➆ mice, 3 times/w for 8 w. Fecal samples were collected from group ➀ and ➁ mice for gut microbiota metagenomic analysis. All mice were sacrificed and mammary, hepatic, muscular and adipose tissues were collected for determining the levels of induced gene expression.

### 1.2 Quantitative polymerase chain reaction (Q-PCR)

RNA isolation, purity determination, electrophoresis monitoring, reverse transcription, and quantification were performed obeying a standard protocol. The relative copy numbers = 2^−ΔΔCт^, in which ΔCт = Ct _target gene_ - Ct _reference gene_, ΔΔCт = ΔC _treatment sample_ - ΔC _control sample_. The primers were designed and applied to amplification under the following amplification condition: 95°C, 30s; 95°C, 3s, 64°C, 34s, 45 cycles. The primers were listed as below:

RUNX1-F: CTGCCCATCGCTTTCAAGGT
RUNX1-R: GCCGAGTAGTTTTCATCATTGCC
TP53BP1-F: ATGGACCCTACTGGAAGTCAG
TP53BP1-R: TTTCTTTGTGCGTCTGGAGATT
BCL11A-F: ACAAACGGAAACAATGCAATGG
BCL11A-R: TTTCATCTCGATTGGTGAAGGG
GAPDH-F: ACAGTCAGCCGCATCTTC
GAPDH-R: CTCCGACCTTCACCTTCC

### 1.3 Enzyme-linked immunosorbent assay (ELISA)

Mouse LPS ELISA Kit was purchased from Jinma (Shanghai, China). Mouse TNF-α and TNF receptor-1 (TNFR1) ELISA Kit was purchased from Chenglin (Beijing, China).

### 1.4 Metagenomic analysis

The gut microbiota profiles in mouse fecal samples were identified by the high-throughput 16S VX sequencing-based classification procedure. The sequencing (sample preparation, DNA extraction and detection, amplicon purification, library construction and online sequencing) and data analysis (paired end-reads assembly and quality control, operational taxonomic units cluster and species annotation, alpha diversity and beta diversity) were conducted by Novogene, Beijing, China.

### 1.5 Statistical analysis

The software SPSS 22.0 was employed to analyze data, and the software GraphPad Prism 5.0 was employed to plot graphs. The Independent Simple Test was used to compare all groups, but the Kruskal-Wallis Test followed by Nemenyi test was used when the data distribution is skewed. The significance level (p value) was set at <0.05 (*), <0.01 (**), <0.001 (***) and <0.0001 (****).

## 2 Results and Discussion

### 2.1 CS promotes SSB overgrowth

In the gut of healthy Chinese people, SSB such as *Bacteroides thetaiotaomicron* J1 and 82 strains, *B. ovitus* E3 strain, and *Clostridium hathewayi* R4 strain were classified^[16]^. Meanwhile, *Akkermansia muciniphila* was also confirmed to be a gut commensal species capable of secreting sulfatases^[17]^. After fed mice with CS, we found the increases in *A. muciniphila* and *B.cereus* abundance, in which *A. muciniphila* accounts for 4% among *Akkermansia*, and 1.059% among all bacterial phyla; *B.cereus* accounts for 4.41% among *Bacillus*, and 1.028% among all bacterial phyla. In contrast, SSB were almost not detected in the gut of control mice, AL1 and AL2. Additionally, *Escherichia coli* producing LPS accounts for 1.62% in *Escherichia*, and 0.379% among all bacterial phyla (**Figure 1**).

**Fig.1.**
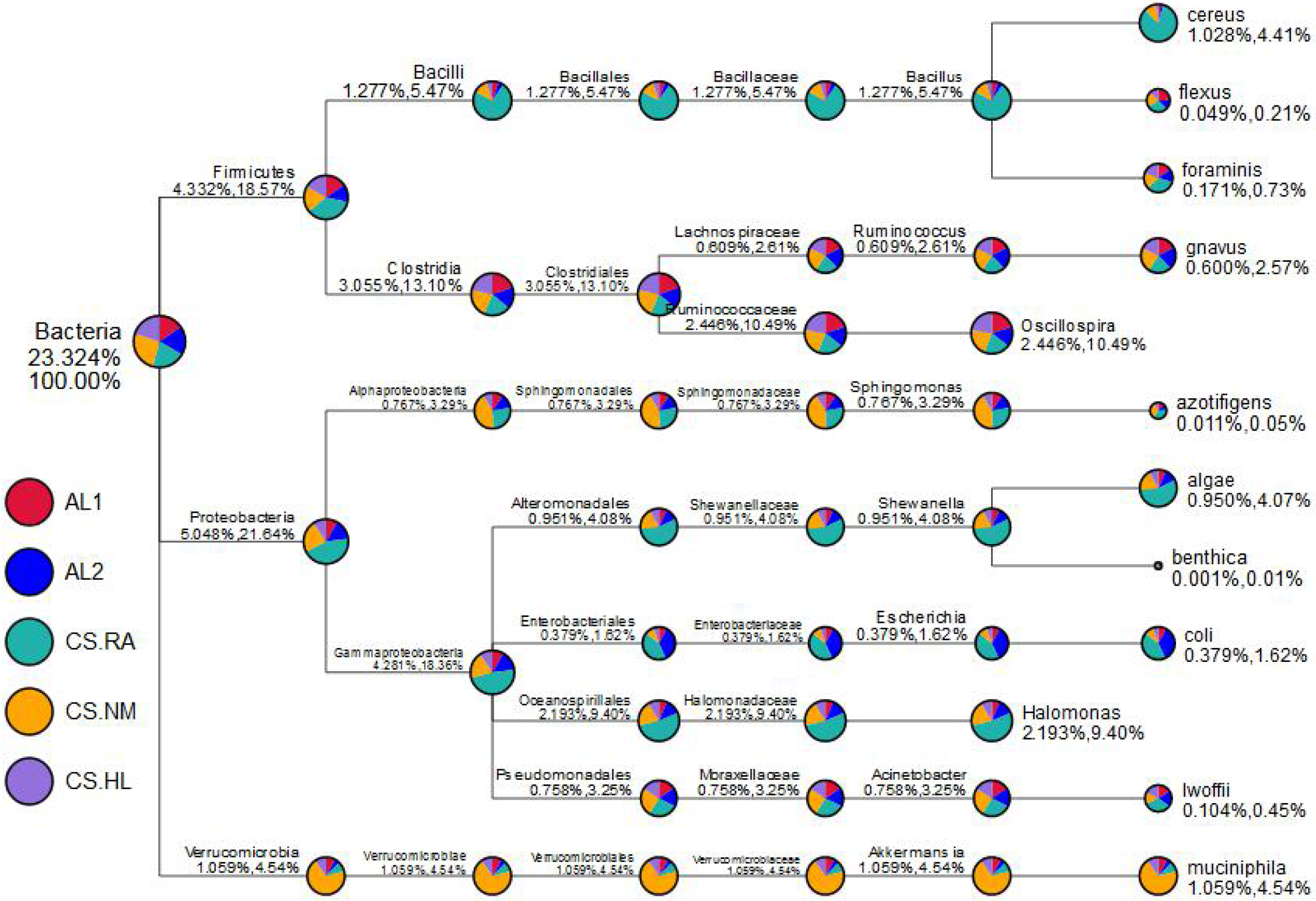
A taxonomic tree of bacteria specific in mouse fecal samples. The fan-shaped sectors in the circle represent different samples corresponding to the left legends, and the proportion of a fan-shaped sector indicates a relative abundance of the sample in the taxonomy. For two percentages listed below the taxonomic nomenclature, the former is the percentage among all species, and the letter is the percentage in the selected species. AL1, AL2: Mice fed *Ad libitum* chow; CS.RA, CS.NM, CS.HL: Mice fed CS.

Although fed CS without exception, the kind and abundance of SSB in the gut of each mouse show the significant variations among individuals. The richness of *A. muciniphila* in the treated mouse, CS.NM, reaches 4.00%, but that in the treated mice, CS.RA and CS.HL, is only 0.50%, which is slightly higher than the average percentages, 0.30%, in the control mice, AL1 and AL2. In contrast, the richness of *B.cereus* in the treated mouse, CS.RA, reaches 4.03%, but that in the treated mice, CS.NM and CS.HL is only 0.24% and 0.65%, which are equal or slightly higher than the average percentages, 0.24%, in the control mice, AL1 and AL2 (**Table 1**).

**Table 1.**
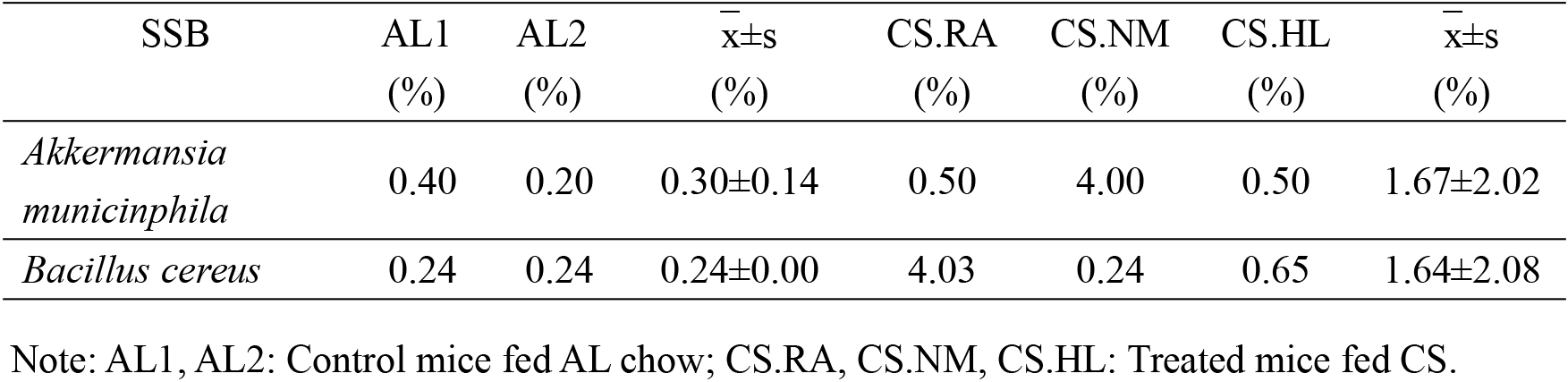
The classification and proportion of gut SSB in mice fed CS

| SSB | AL1 | AL2 | $\bar{x}\pm s$ | CS.RA | CS.NM | CS.HL | $\bar{x}\pm s$ |
| --- | --- | --- | --- | --- | --- | --- | --- |
|  | (%) | (%) | (%) | (%) | (%) | (%) | (%) |
| *Akkermansia muciniphila* | 0.40 | 0.20 | 0.30±0.14 | 0.50 | 4.00 | 0.50 | 1.67±2.02 |
| *Bacillus cereus* | 0.24 | 0.24 | 0.24±0.00 | 4.03 | 0.24 | 0.65 | 1.64±2.08 |

Note: AL1, AL2: Control mice fed AL chow; CS.RA, CS.NM, CS.HL: Treated mice fed CS.

These above results indicated that feeding CS to mice can change the gut microbiota composition, but not every mouse exhibits opportunistic infection by SSB, implying that the remarkable difference of SSB occurs among the gut of each individual mouse. Mounting evidence shows that the effect of CS on the female mice are more predominately than the male mice^[18]^. Our research has chosen the female mice for investigation, so the great individual variation of gut microbiota communities can be attributed to CS.

### 2.2 SSB prompts serum LPS and pro-inflammatory mediator fluctuation

To explore whether CS feeding would induce inflammatory responses by leaking gut microbiota debris, we determined the serum LPS and TNF-α levels in mice, as listed in **Table 2**.

**Table 2.** The individual specificity of serum LPS and TNF-α in mice fed CS

| Serum titer | AL1 | AL2 | $\bar{x}\pm s$ | CS.RA | CS.NM | CS.HL | $\bar{x}\pm s$ |
| --- | --- | --- | --- | --- | --- | --- | --- |
| LPS (EU/L) | 11.96 | 14.94 | 13.45±2.11 | 16.18 | 16.09 | 13.29 | 15.19±1.64 |
| TNF-α<br>(ng/L) | 378.73 | 526.25 | 452.49±104.31 | 1067.18 | 407.87 | 664.67 | 820.69±298.55 |

Note: AL1, AL2: Control mice fed AL chow; CS.RA, CS.NM, CS.HL: Treated mice fed CS.

From the above results, it was clear that CS feeding-induced gut opportunistic infection increases the serum LPS and TNF-α levels albeit no significant statistical difference between AL mice and CS mice. This phenomenon seems to be resulted from the huge individual variation of the compared raw data.

The tissue variations in LPS, TNF-α and TNFR1 levels were also observed. In AL1 and CS.RA mice, for example, LPS levels are declined in the hepatic, adipose and muscular tissues, whereas TNF-α levels are declined only in the adipose tissue, but are elevated in the adipose and muscular tissues. TNFR1 levels are elevated in the adipose and muscular tissues (**Figure 2**), indicating a conversion of LPS from high levels to low levels.

**Fig.2.**
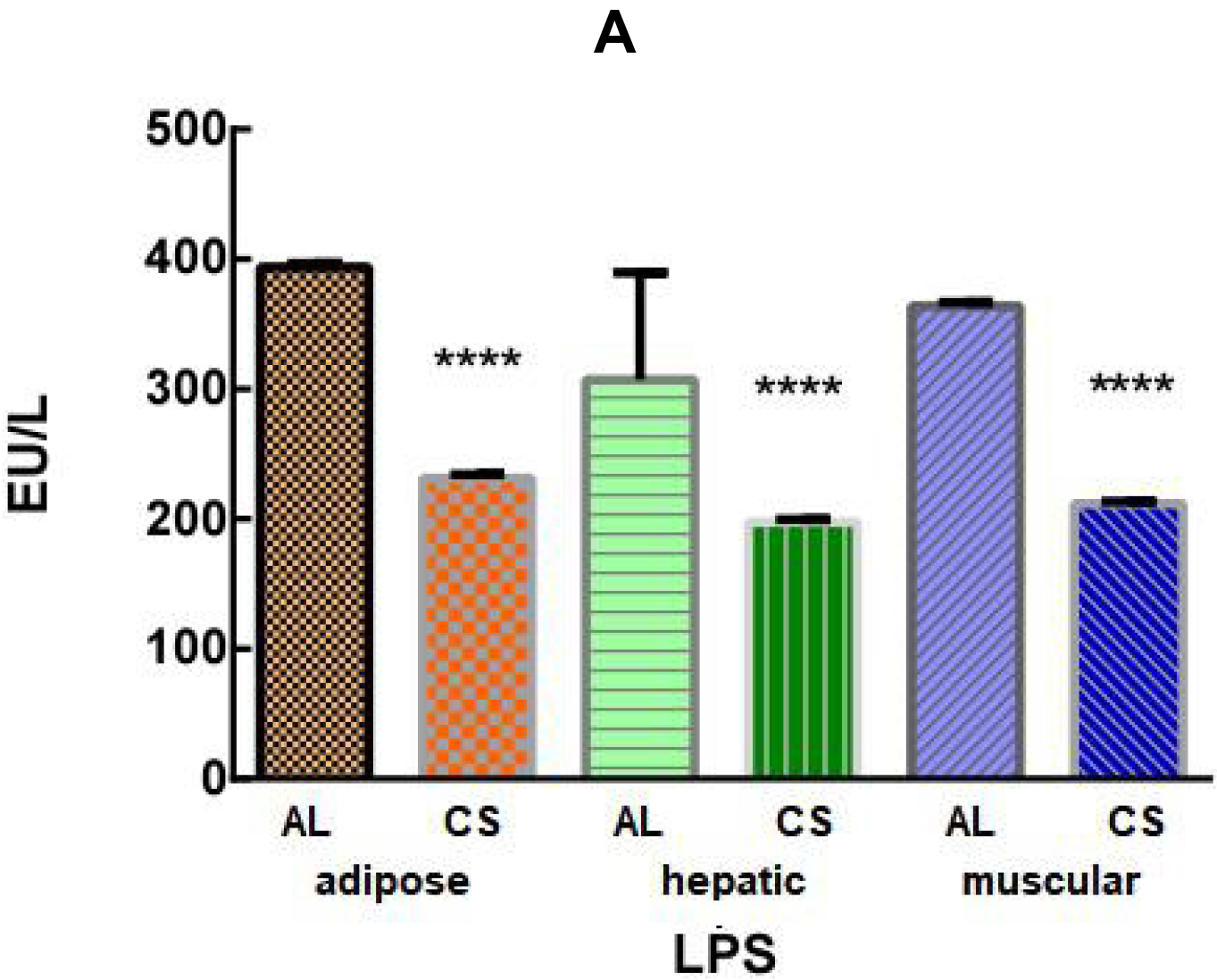
The tissue specificity of LPS and pro-inflammatory mediators in mice fed CS. A. LPS levels in mouse liver, adipose, and muscular tissues; B. TNF-α levels in mouse liver, adipose, and muscular tissues; C. TNFR1 levels in adipose and muscular tissues. AL: Tissues from mice fed *Ad libitum* chow; CS: Tissues from CS-treated mice. **** *p*<0.0001 (*n*=15).

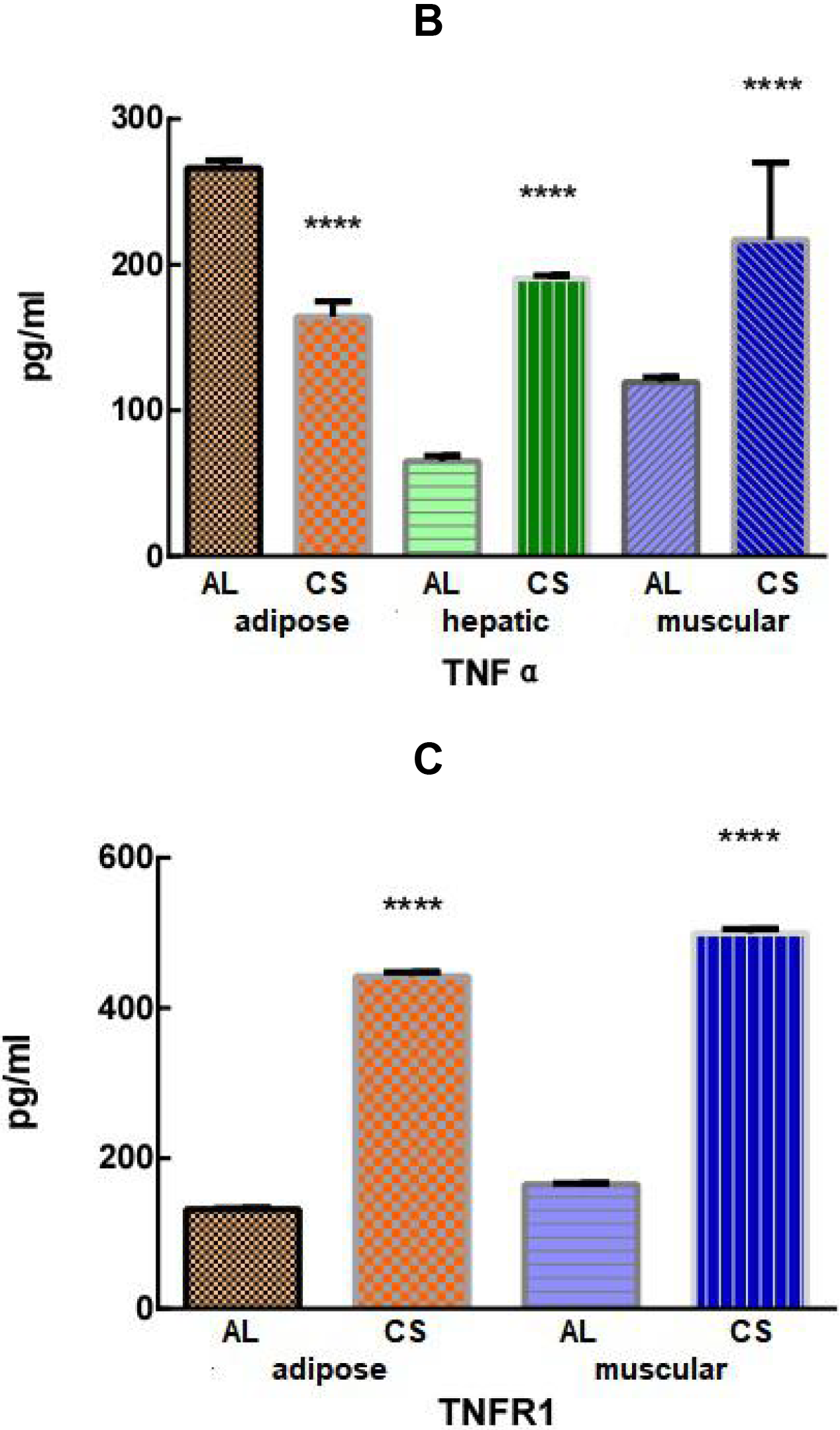

The tissue variations of LPS and pro-inflammatory cytokines/receptors may be associated with the serum and tissue LPS levels and anti-LPS depleting LPS rates. Hersoug et al^[19]^ disclosed that after scavenger receptor B1 (SRB1) binds to LPS, SRB1-LPS can then be incorporated into the chylomicrons, which enhances LPS trespassing across the endothelium and entering into adipocytes. These results implied that LPS may be preferably transported into the adipose tissue, which induces anti-LPS and upregulates TNF-α and TNFR1, eventually eradicating LPS and downregulating TNF-α.

### 2.3 Induced expression of mouse mammary gland tumor-related transcription factor genes

From the data of metagenomic analysis, it was predicted that the individual variation of mouse gut microbiota difference may represent the major cause of distinct inflammatory severity. So when we fed CS, we also gastrically administered BC, by which the individual variation of mouse gut microbiota difference was anticipated to be minimized. After CS+BC feeding or LPS injection, we quantified the mouse mammary mRNA levels of the mammary tumor-related transcription factor genes *BCL11A*^[20]^ and *RUNX1*^[21]^, and the tumor suppressor TP53 binding protein-1 gene (*TP53BP1*)^[22]^.

The results indicated that *BCL11A* and *RUNX1* mRNA levels are higher in CS+BC or LPS-treated mice than those in control mice, in which LPS-induced mRNA levels are higher than CS+BC-induced mRNA levels. As to *TP53BP1* mRNA, both treatments induce the similar levels with control. Besides, CS+BC+FS significantly increase *RUNX1* mRNA (**Figure 3**).

**Fig. 3.**
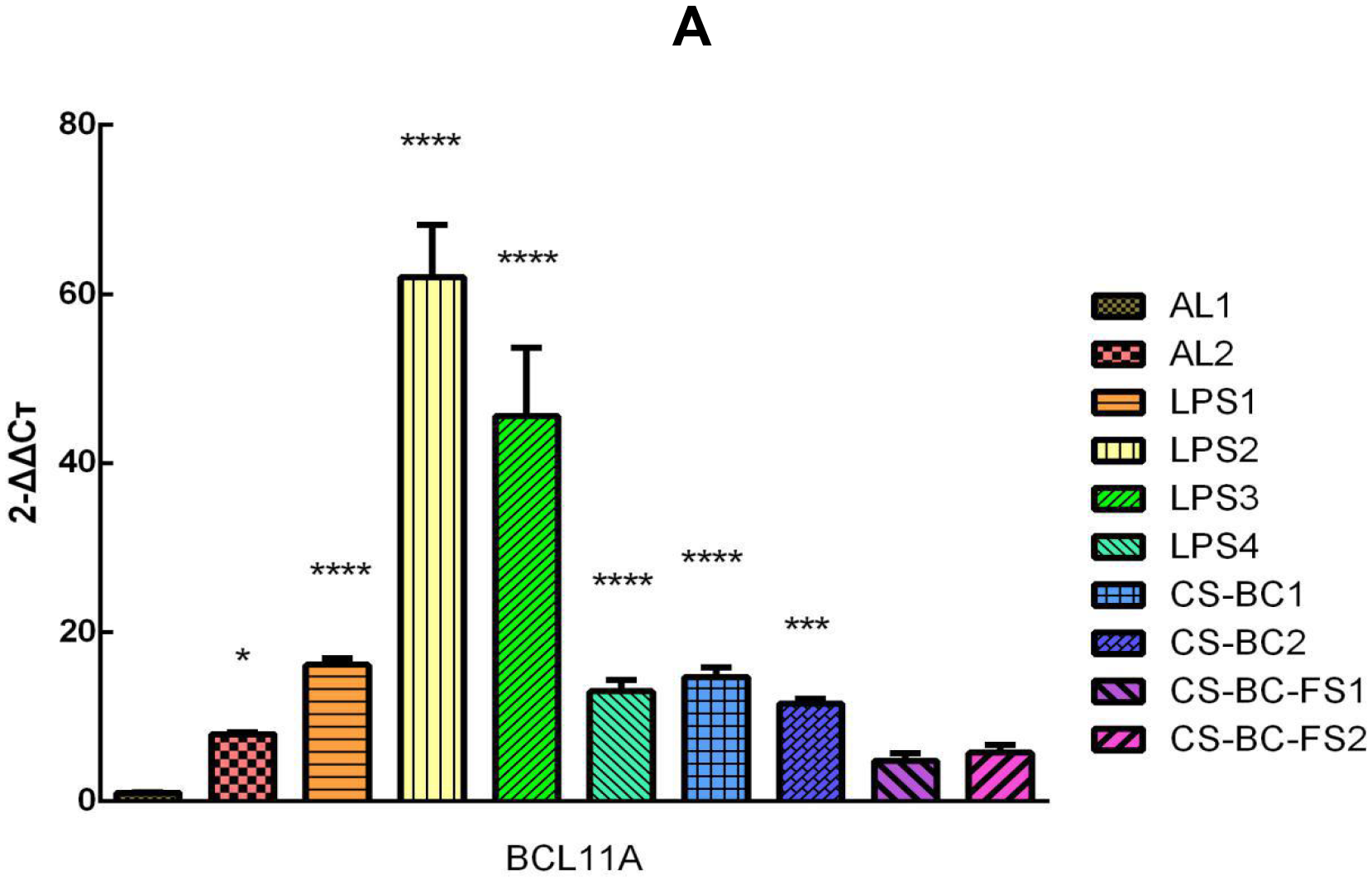
The multi-factorial induction of tumor-related genes expression in mouse mammary tissues. A. The relative *BCL11A* mRNA copy numbers; B. The relative *RUNX1* mRNA copy numbers mRNA; C. The relative *TP53BP1* mRNA copy numbers. AL: Mammary gland tissues of mice fed *Ad libitum* chow; LPS: LPS-treated mouse mammary gland tissues; CS+BC: chondroitin sulfate+*Bacillus cereus*-treated mouse mammary gland tissues; CS+BC+FS: chondroitin sulfate+*Bacillus cereus*+fulvestrant-treated mouse mammary gland tissues. \**p*<0.05; \*\**p*<0.01; *** *p*<0.001; **** *p*<0.0001 (*n*=15).

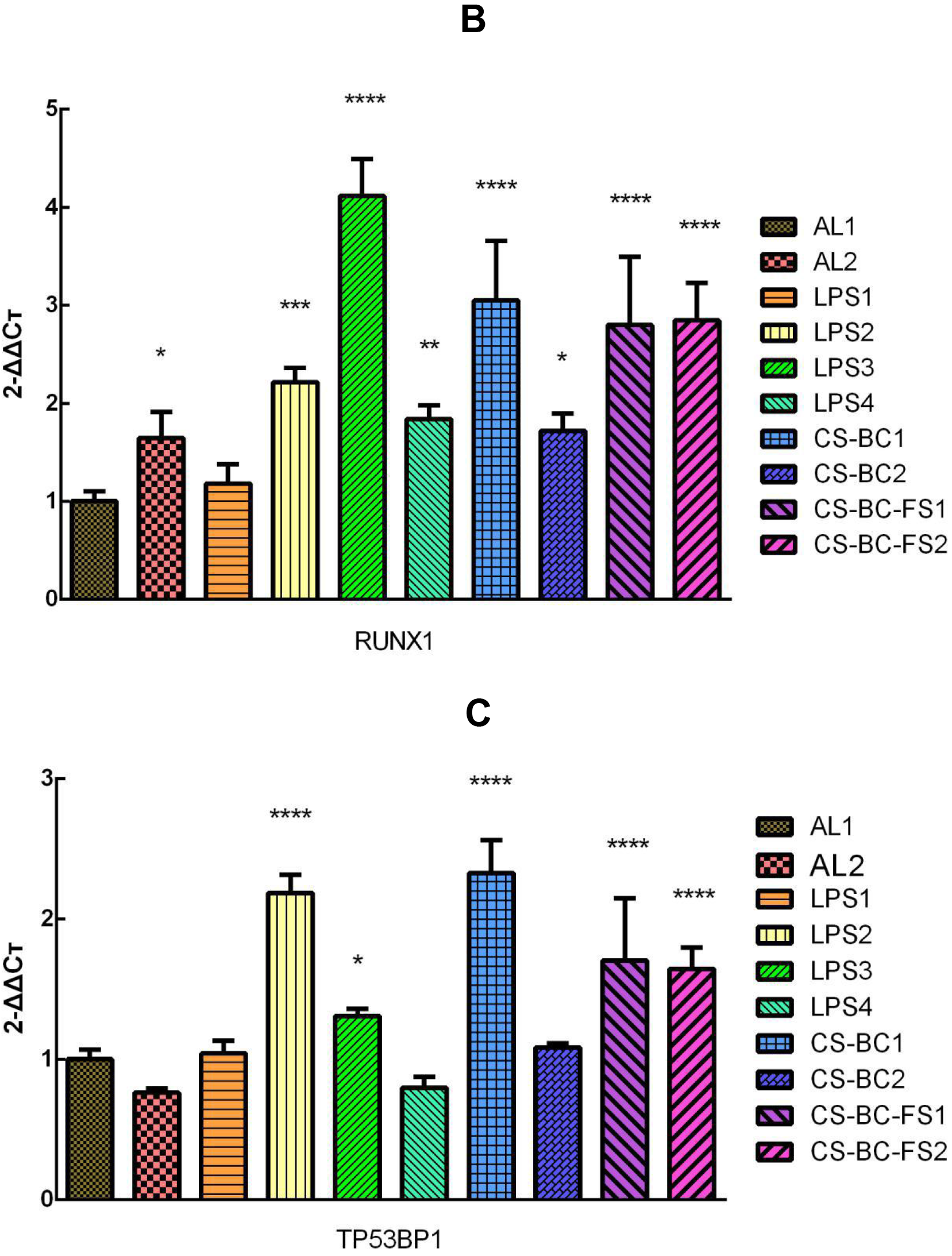

It could be seen from the above figure that the expression levels of tumor-related genes remain extraordinarily varied, reflecting that the attempt of uniforming the gut microbiota using BC, which is perhaps because the activity of *in vitro* cultured bacteria is relatively low, and the orally administered bacteria are killed by the acidic gastric juice.

### 2.4 Exogenous estradiol fails to induce *BCL11A*, *RUNX1* and *TP53BP1* expression

To evaluate the effect of ED on mammary tumor incidence, we quantified the mammary levels of *BCL11A*, *RUNX1* and *TP53BP1* mRNA in ED-injected mice. The results shown that the mammary *BCL11A*, *RUNX1* and *TP53BP1* mRNA levels induced by either high-level ED (4.0 mg/kg) or low-level ED (0.5 mg/kg) are comparative to those in control mice (**Figure 4**).

**Fig. 4.**
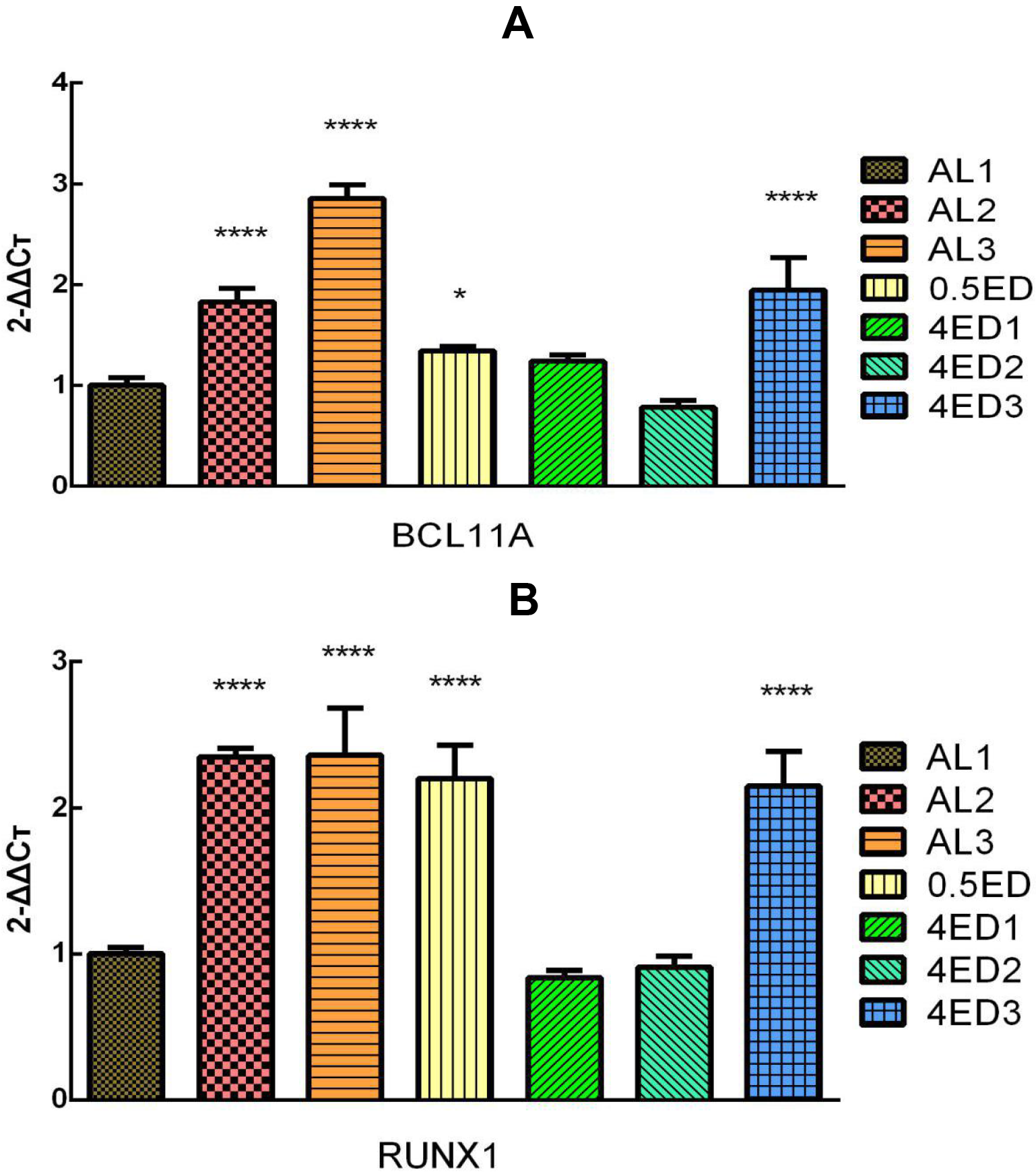
The estradiol-induced expression of tumor-related genes in mouse mammary gland tissues. A. The relative *BCL11A* mRNA copy numbers; B. The relative *RUNX1* mRNA copy numbers mRNA; C. The relative *TP53BP1* mRNA copy numbers. AL: Mammary gland tissues of mice fed *Ad libitum* chow; 0.5ED: 0.5mg/kg estradiol-treated mouse mammary gland tissues; 4ED: 4mg/kg estradiol-treated mouse mammary gland tissues. \**p*<0.05; \*\**p*<0.01; **** *p*<0.0001 (NC: *n*=9; 0.5E: *n*=3; 4E: *n*=9).

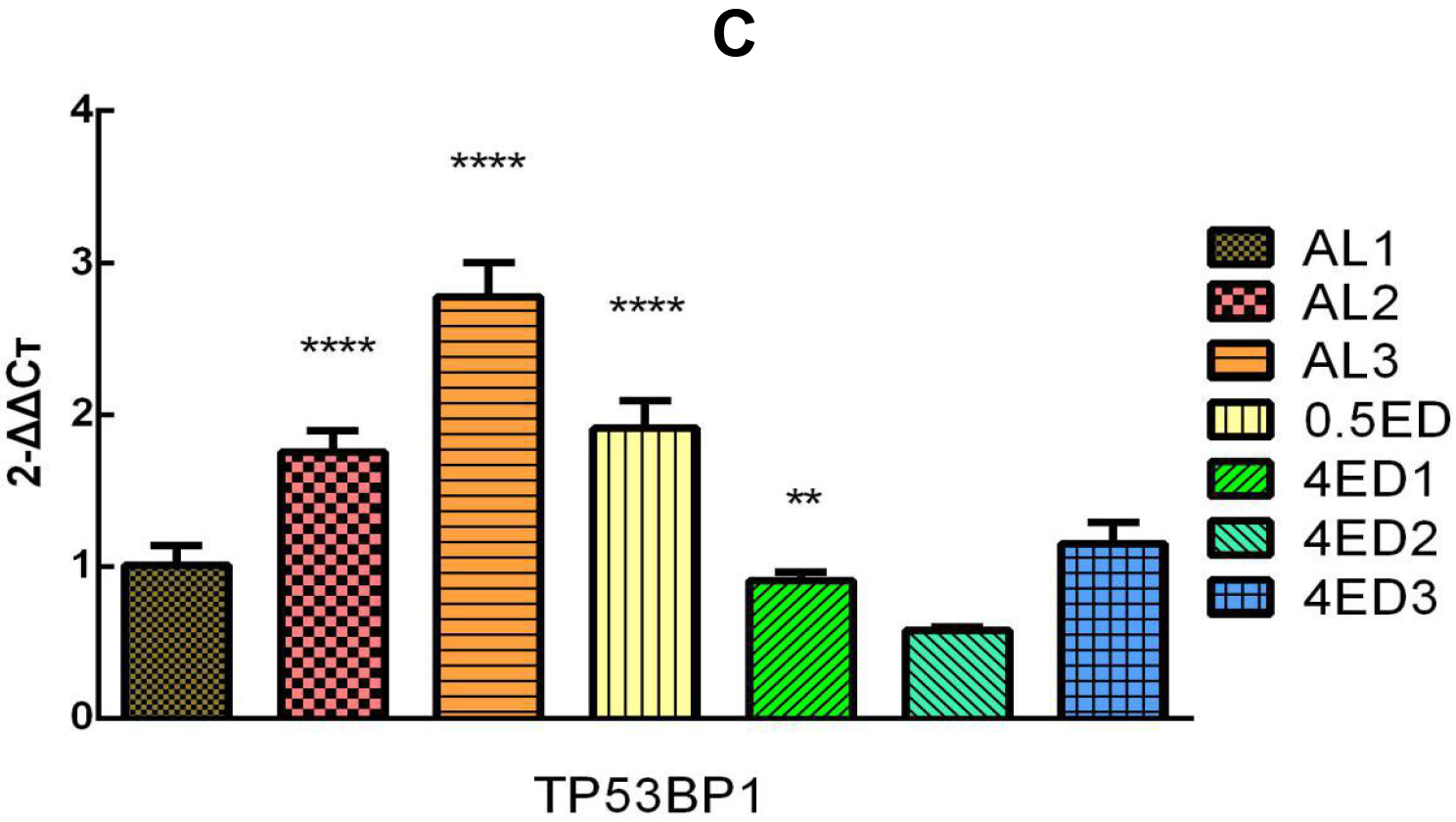

Accumulating evidence shows that the incidence of breast cancer among women in the industrialized countries is most probably related to the environmental contamination by estrogen-like chemicals^[23]^. Nevertheless, it has been unknown how can estrogens *per se* trigger breast cancer. From the carcinogenic mechanism, however, it has been clear that estradiol can produce 2-hydoxyestradiol and 4-hydoxyestradiol at first, and then produce semi-quinone and quinone, which can react with DNA to form depurinated nucleotides and convert A-T to GC (point mutation)^[13]^. Additionally, estrogens can also increase the tumorigenic risk by inducing the methylation of repetitive DNA sequences, such as LINE-1 and Alu family^[24]^.

### 2.5 Induced expression of mouse mammary gland tumor marker genes

By quantifying the expression levels of mammary tumor marker genes, we found that CS+BC, like LPS, can decline the levels of mammary myoepithelium marker CK5-6^[25]^, but elevate the levels of mammary hyperplasia marker Ki-67^[26]^. As to the tumor metastasis marker E-cadherin^[27]^ and the tumor differentiation marker GCDFP-15^[28]^, no distinguishable changes were noted, suggesting that mammary tumor has been just formed, but has not yet entered the stage of metastasis and differentiation (**Figure 5**).

**Fig. 5.**
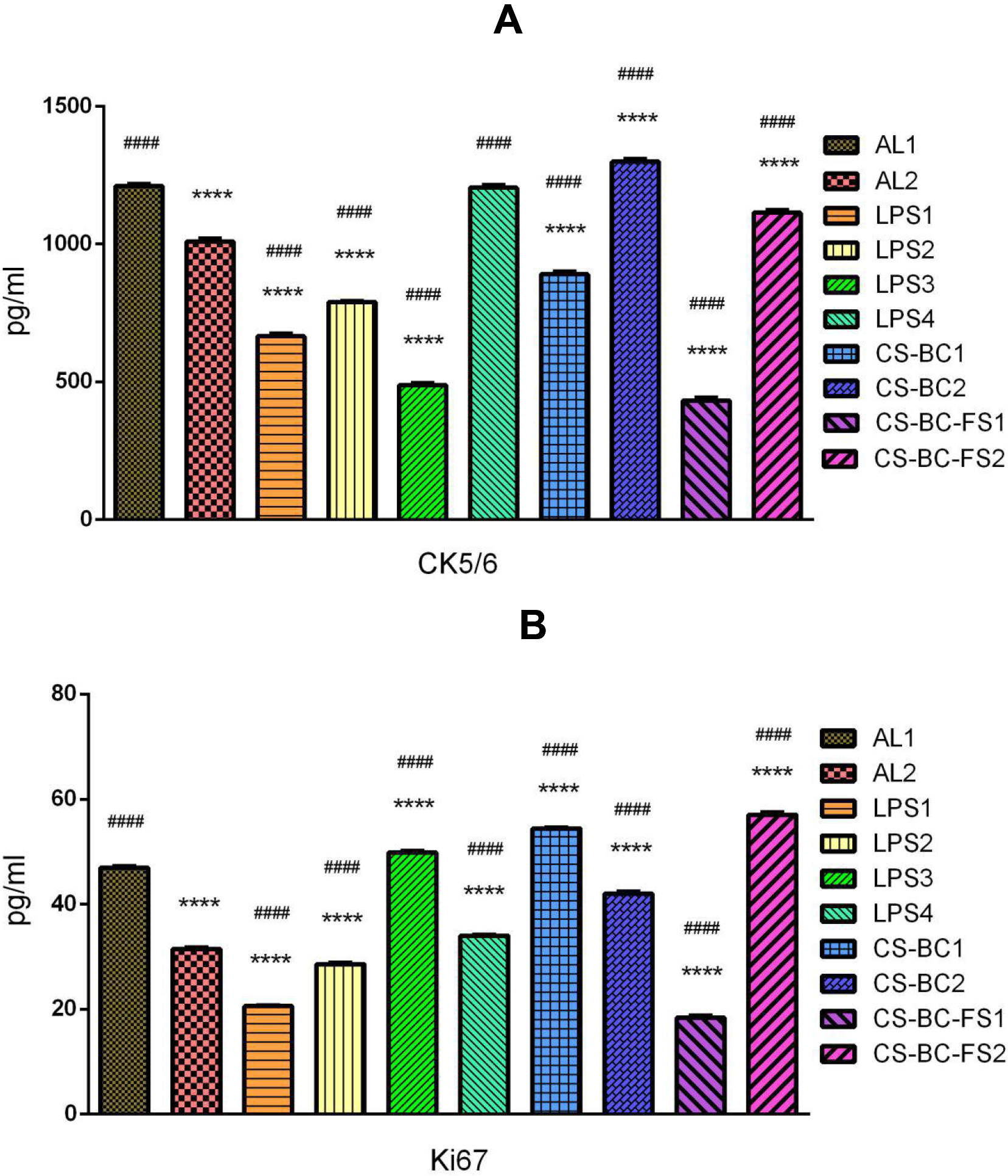
The multi-factorial induction of tumor marker expression in mouse mammary gland tissues. A. CK5-6 levels; B. KI67 levels; C. E-cadherin levels; D. GCDFP-15 levels. AL: Mammary gland tissues of mice fed *Ad libitum* chow; LPS: LPS-treated mouse mammary gland tissues; CS+BC: chondroitin sulfate+*Bacillus cereus*-treated mouse mammary gland tissues; CS+BC+FS: chondroitin sulfate+*Bacillus cereus*+fulvestrant-treated mouse mammary gland tissues. * Compared to NC1; ^#^ Compared to NC.*p*<0.05; **or^##^*p*<0.01; ****or^####^*p*<0.0001 (NC: *n*=6; LPS *n*=12; CS+BC: *n*=6; CS+BC+FS: *n*=6).

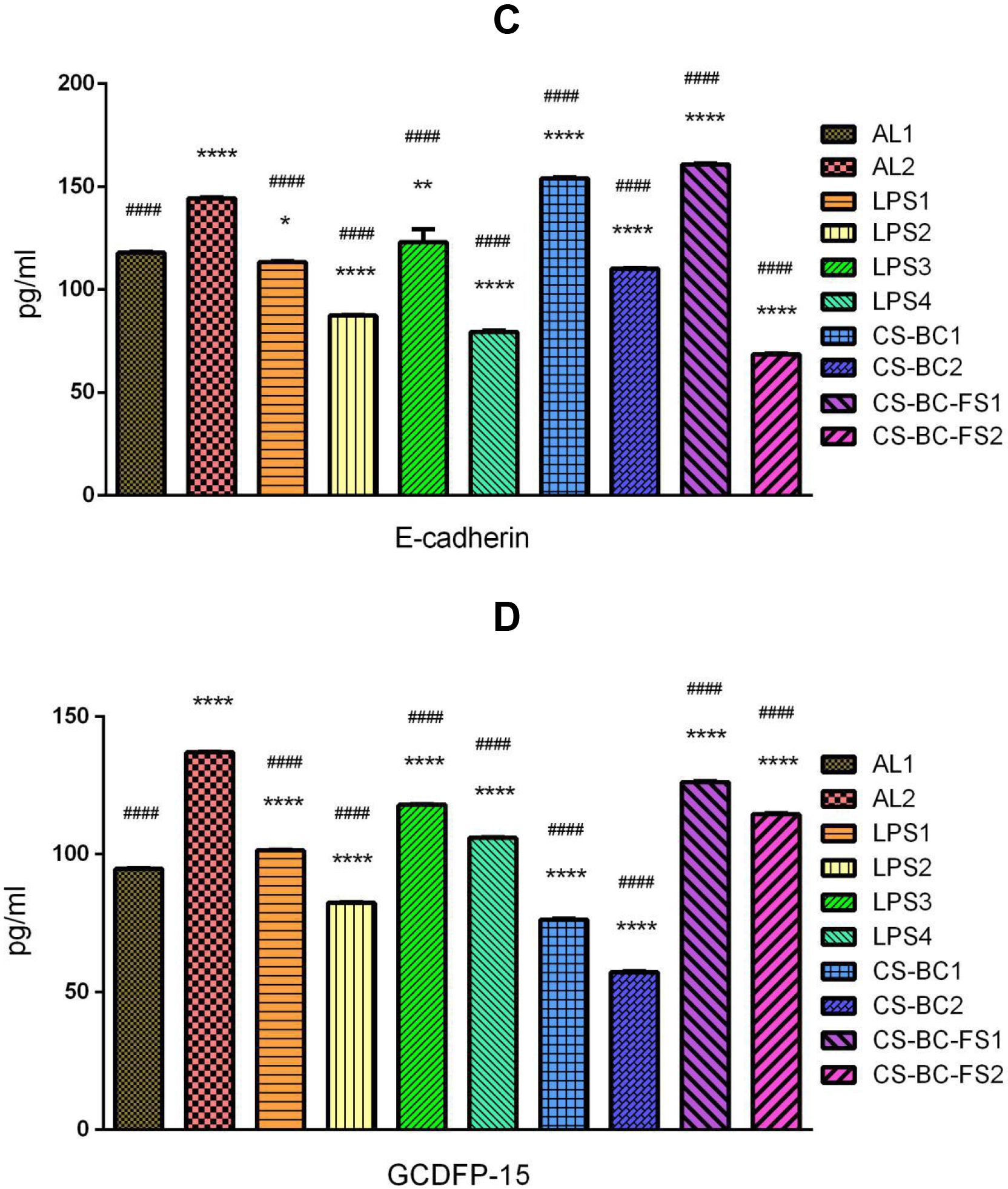

From the above data, the individual variation remains a typical character, reflecting an affection from the individual gut microbiota variation. In similar, the expression levels of tumor marker genes are also individually variable, in which some are high expressed, while others are low expressed.

### 2.6 Hypoxia-induced expression of angiogenesis genes

To mimic the inflammatory inactivation of ERs, we employed the ER agonist FS to treat CS+BC mice, and determined the serum ED level and hepatic HIF-1α and VEGF levels. As results, it was shown that ED levels are higher in treated mice than those in control mice, addressing that FS can inhibit ER activity and complementarily increase ED levels. It was also observed that HIF-1α levels are unchanged, whereas VEGF levels are dramatically elevated (**Figure 6**).

**Fig. 6.**
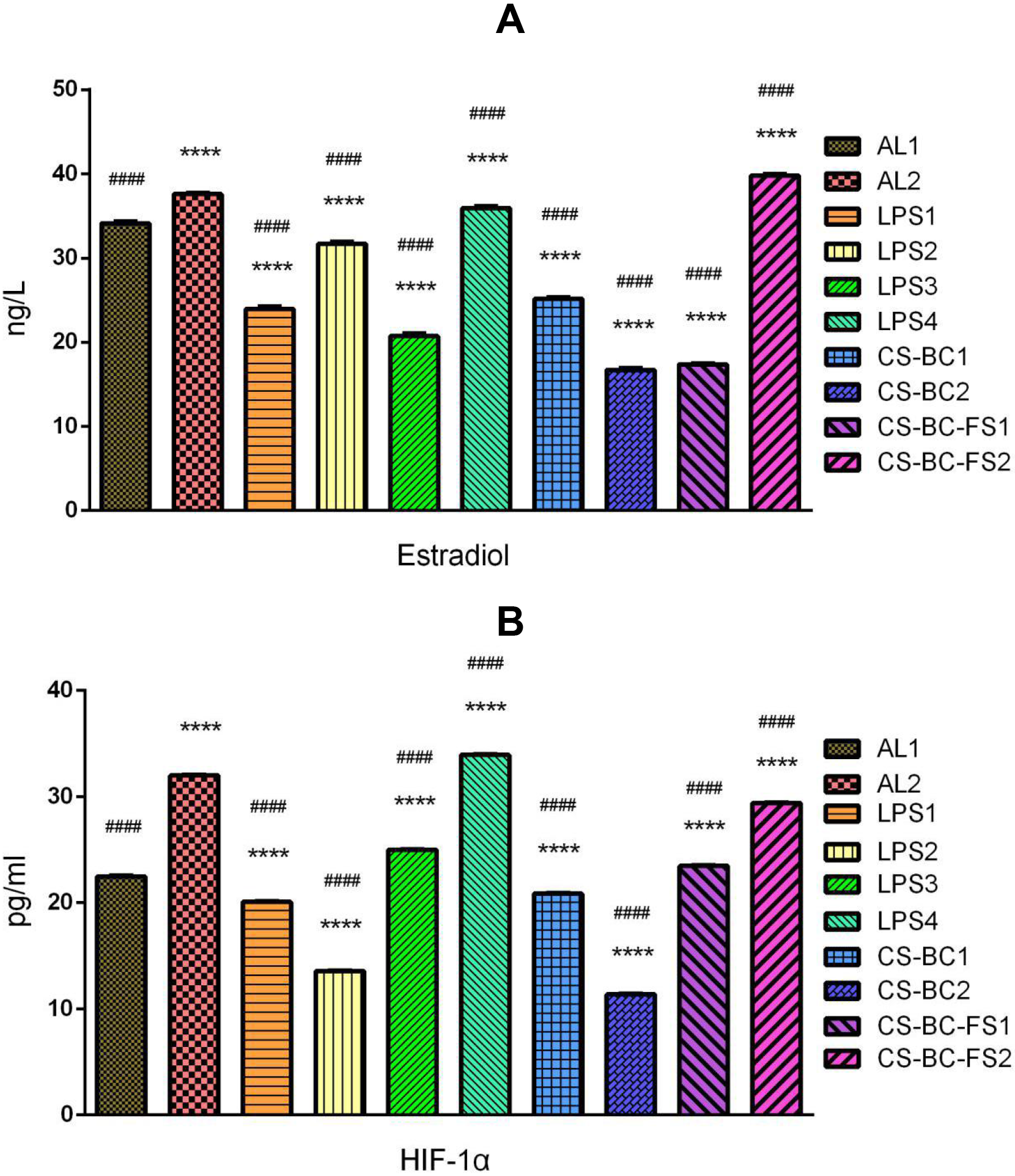
The multi-factorial induction of mouse serum estradiol and liver HIF-1α and VEGF levels. A. Estradiol levels; B. HIF-1α levels; C. VEGF levels. AL: Serum or liver tissues of mice fed *Ad libitum* chow; LPS: LPS-treated mouse serum or liver tissues; CS+BC: chondroitin sulfate+*Bacillus cereus*-treated mouse mouse serum or liver tissues; CS+BC+FS: chondroitin sulfate+*Bacillus cereus*+fulvestrant-treated mouse mouse serum or liver tissues. * Compared to NC1; ^#^ Compared to NC. *p*<0.05; **or^##^*p*<0.01; ****or^####^*p*<0.0001 (NC: *n*=6; LPS: *n*=12; CS+BC: *n*=6; CS+BC+FS: *n*=6).

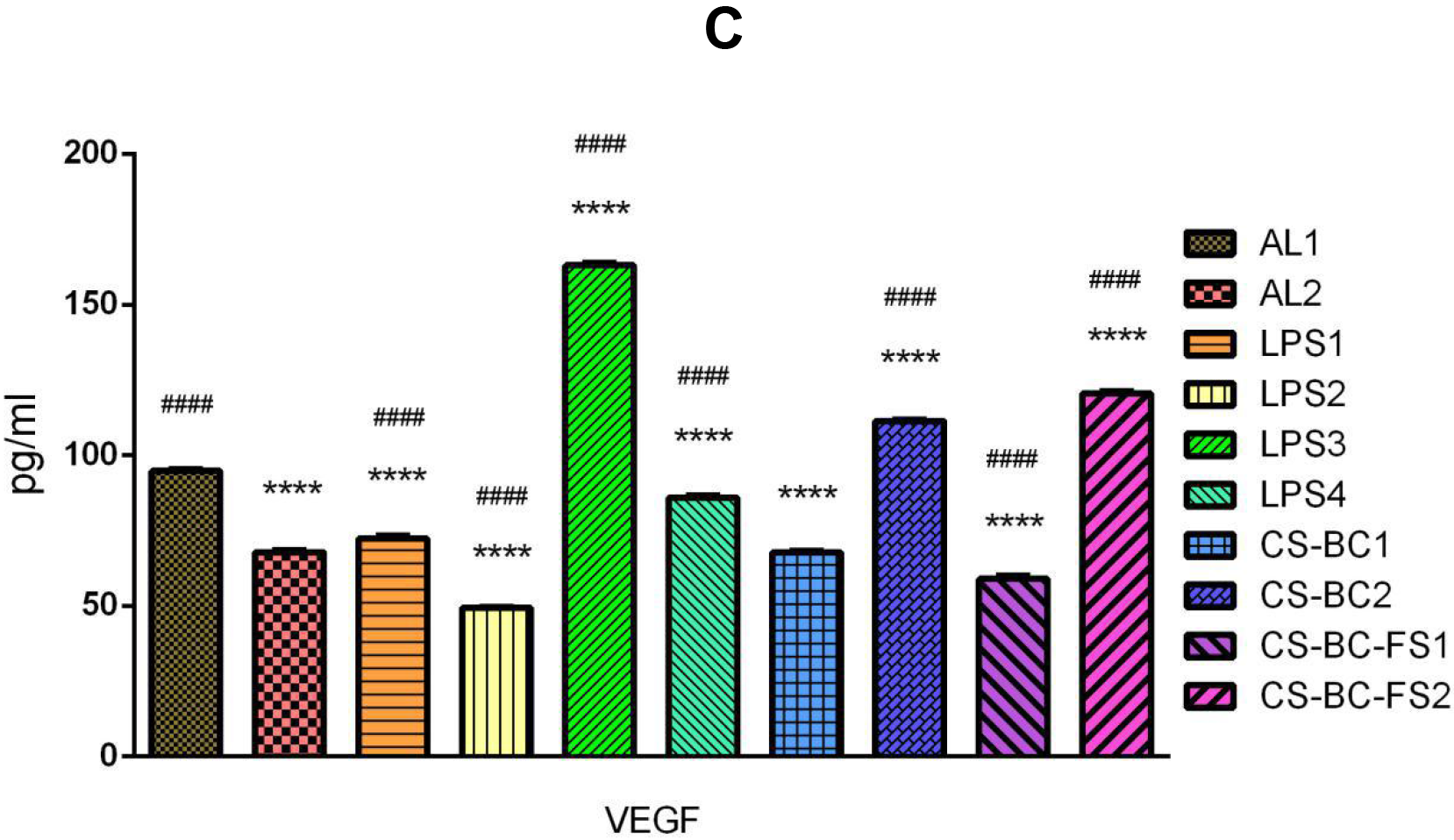

It was known that the proinflammatory cytokine-activated hypoxia can induce VEGF expression and angiogenesis by HIF-1α^[29]^. It was also evident that the expression of HIF-1α is regulated at the post-translational level (activated upon translocation from the cytoplasm to the nuclei) rather than transcriptional and translational levels^[30]^.

## 3 Conclusion

The present study preliminarily testified the hypothesis of gut origin of breast cancer. By CS mimicking a meat diet, BC mimicking gut opportunistic infection, LPS mimicking hyper-endotoxinemia, ED mimicking hyperestrogenemia, and FS mimicking ER inflammatory inactivation, we obtained some evidence regarding the induced gene expression of mammary tumor-related transcription factors and tumor markers, and explained the tumor heterogeneity by the individual and tissue variations of gut dysbiosis and systemic inflammation.

Our next work should be re-acting a whole process of breast cancer at the cellular, tissue and individual levels, and eventually eradicating the risk of breast cancer by maintaining gut homeostasis.

## Acknowledgements

We thank Ms. Fan Pei in Novogene (Beijing, China) for her assistance in meta-genomic analysis of the gut microbiome. This work was supported by the National Science Foundation of China (Grant No. 81273620; 81673861; 81774041).

